# Utilizing raw rapeseed press cake in foods: A case study on sensory quality and profile of selected bitter compounds in snack bars

**DOI:** 10.64898/2026.03.20.712648

**Authors:** Jakob Skytte Thorsen, Andrea Bononad-Olmo, Arendse Maria Toft, Niels Christian Holm Sanden, Kwadwo Gyapong Agyenim-Boateng, Michal Poborsky, Christoph Crocoll, Barbara Ann Halkier, Wender L.P. Bredie, Deyang Xu

## Abstract

Today’s canola quality rapeseed press cake (RPC) is a protein-rich co-product with potential as human food, but its application is limited due to antinutritional compounds and bitter taste. It remains, however, unknown how introduction of raw RPC to a food matrix affects sensory perception and which metabolites drive the sensation. Here, raw RPC from whole or dehulled seeds was introduced into snack bars at 0%, 7%, 14%, and 21%, and sensory responses were correlated to selected known RPC-derived bitter compounds. A trained panel evaluated 13 RPC-characteristic sensory attributes, and the bitter compounds sinapic acid, kaempferol 3-O-(2‴-O-sinapoyl-β-sophoroside) (KSS), KSS-hexose, selected bitter glucosinolates, and goitrin were quantified using targeted LC–MS/MS. Most dose-dependent sensory responses increased up to 14% RPC and then plateaued, whereas astringent mouthfeel increased almost linearly across the full dose range. Dehulling intensified several odor- and flavor-related attributes but did not increase bitterness or protein content in the final product. Principal component analysis linked bitterness and astringency positively with KSS, KSS-hexose, and goitrin. Dose-over-threshold analysis further showed that goitrin, but not progoitrin, reached concentrations relevant for bitterness perception. Together, the results demonstrate that raw RPC contributes distinct dose-dependent sensory attributes and that metabolite transformations in the food matrix shape final sensory profiles. These findings provide a basis for developing RPC-containing foods and for breeding rapeseed lines with improved sensory characteristics.

**HIGHLIGHTS:**

- This study presents the first sensory panel assessment of rapeseed press cake (RPC)-containing in food products (snack bars) made from whole and dehulled seeds.
- 13 RPC-characteristic sensory attributes are identified.
- Sensory profiles of the tasted snack bars differed significantly, influenced by the dosage of RPC and by the dehulling treatment. Bitterness and astringency are positively correlated with the RPC dosage.
- Goitrin, kaempferol 3-O-(2‴-O-sinapoyl-β-sophoroside) (KSS) and sinapic acid are RPC-derived bitter compounds that correlate with bitter taste of RPC-containing snack bars.
- Approximately 90% of glucosinolates introduced with the RPC are not detected in the snack bars, and goitrin levels in snack bars accounts for only ∼10% of introduced progoitrin.
- Goitrin is – for the first time – reported to contribute to the perceived bitterness of an RPC-containing food product.

## 1. INTRODUCTION

The Planetary Health Diet calls for a substantial shift from animal- to plant-based protein foods to achieve benefits both for human and planetary health (Willett et al., 2019). Rapeseed (*Brassica napus L.*) is the world’s second-largest oilseed crop and the rapeseed press cake (RPC) that remains after oil extraction is a protein-rich (30-40%) side stream that today is primarily used as feed and largely unexploited as protein source for human consumption (Nwokolo & Bragg, 1978; Wia̧z et al., 2005). Application of raw, unprocessed RPC from today’s double low elite lines in foods faces two challenges. Firstly, the raw RPC is not approved as food in EU due to high levels of anti-nutritional glucosinolates (EFSA NDA Panel, 2023; Thorsen et al., 2025). Secondly, consumers may regard RPC as feed with undesirable bitterness as evidenced in a Polish study, in which consumers tasted biscuits containing up to 20% RPC (Szydłowska-Czerniak et al., 2021).

Key metabolites associated with bitterness of seeds from rapeseed have been identified. This includes the phenolic sinapine and sinapic acid, the former present at ∼1-2% of rapeseed seeds (Ismail et al., 1981; Schwarze, 1949). Schwarze (1949) reported that sinapine (the choline ester derivative of sinapic acid) was key for bitterness of raw RPC. This was later confirmed by Ismail et al. (1981), who reported that sinapine was accountable for ∼37-50% of the observed bitterness in water slurries of RPC, and that not only sinapine but also its two precursors (sinapic acid and choline chloride) were perceived as bitter. More recently, Hald et al. (2019) identified kaempferol glycosides e.g. kaempferol 3-O-(2‴-O-sinapoyl-β-sophoroside) (KSS) as the major contributor of bitterness in a rapeseed protein isolate. Walser et al. (2024) subsequently identified additional bitter KSS derivatives, although KSS itself remained the primary contributor to the bitterness of the protein isolate. Some glucosinolates (progoitrin, gluconapin, glucobrassicin) have been reported to have bitter taste (Bell et al., 2018). A study on Brussels sprouts reported that progoitrin (2-(R)-OH-3-butenyl glucosinolate), a major glucosinolate in rapeseed seeds, tastes bitter and that its hydrolysis product goitrin (L-5-vinyl-2-thiooxazolidone) tastes extremely bitter (Fenwick et al., 1983).

Rather than using highly processed protein isolates for human consumption, it would be desirable from a health and sustainability perspective to use unprocessed (raw) RPC (Dicken et al., 2025; Lie-Piang et al., 2021). Upon preparation of food products containing RPC, metabolites and enzymes from different ingredients are brought together in a food matrix, which may result in notable changes to the metabolite composition, and thus sensorially alter the final food product compared to that of the individual ingredients. Generally, we lack knowledge about how the different RPC-derived bitter compounds affect sensory quality when in a food matrix.

Towards development of RPC-containing food products, it is important to identify sensory and metabolic bottlenecks in consumer acceptance. Thus, we conducted a sensory assessment with a trained panel to characterize how dose and dehulling treatment impact the intensity of rapeseed-derived sensory attributes in a food matrix. We formulated whole food snack bars with increasing proportion of raw RPC (from 0-21%) as food prototype and combined sensory evaluation with targeted metabolite analysis of known bitter compounds. Here, we report how the effect of increasing the proportion of raw RPC in whole food snack bars affect their sensory quality, and the perceived bitterness is aligned with the presence of key RPC-derived bitter compounds. The correlation between sensory perception and bitter compounds is discussed.

## 2. Materials and methods

### 2.1. Production of rapeseed press cake (RPC)

Both whole and dehulled ‘canola quality’ seeds (<25 µmol/g glucosinolates and <2% eruca acid) were provided by the commercial oil mill ‘Bornholm’s Oliemølle A/S’, Aakirkeby, Denmark. For the dehulled seeds, remaining hulls were removed with an air blower. The whole and dehulled seeds were ‘cold-pressed’ (after pre-heating, no external heat was added, temperature reported by instrument outside press head between 70-80°C during pressing) in a OW 100 s-inox oil press from Ölwerk GmbH. The freshly produced RPC was stored in an air-tight container at 5°C until preparation of snack bars.

### 2.2. Production of snack bars

For the main experiment, seven types of experimental snack bars were produced with increasing proportions of raw RPC, from respectively, whole and dehulled seeds by the Copenhagen-based company ‘Spora’. The base paste consisted of 33 g roasted almonds, 33 g brewed filter coffee, 33 g flax seed, 33 g pumpkin seed, 17 g chia seeds, 117 g Medjool dates and 83 g raisins. Aliquots of the base were proportionally substituted with RPC from cold-pressed whole or dehulled seeds to contain 7, 14 and 21% RPC (**Table 1**). The different pastes were moulded into rectangle bars of 10 g. The bars were packed in airtight containers and stored at 4°C until sensory analysis or were frozen (-20°C) until metabolite analysis.

**Table 1:** Overview of different snack bars used in study. Snack bars contained increasing content (0-21%) of rapeseed press cake (RPC) from either whole or dehulled seeds.

| Sample code | % RPC (w/w) | RPC type |
| --- | --- | --- |
| Control | 0 | None |
| Whole_7 | 7 | Whole |
| Whole_14 | 14 | Whole |
| Whole_21 | 21 | Whole |
| Dehulled_7 | 7 | Dehulled |
| Dehulled_14 | 14 | Dehulled |
| Dehulled_21 | 21 | Dehulled |

### 2.3. Sensory descriptive analysis

#### 2.3.1. Sensory panel

Ten participants were recruited from the external trained sensory panel of the Department of Food Science at the University of Copenhagen. Inclusion criteria were that participants were between 18 and 60 years of age, have no known food allergies, not to be pregnant or lactating, and could understand and communicate in English. Each subject participated by informed consent.

#### 2.3.2. Descriptive sensory profiling

The trained panellists received a two-hour training session, which included vocabulary development, attribute definition, reference selection and ranking of seven samples. A subsequent training session included similar activities, refining attributes and intensity scaling of samples. Training was deemed sufficient, as no significant interaction effects between replicates and assessors were observed.

The descriptive sensory profiling was conducted in sensory booths designed according to ISO/ASTM standards. Portions of 10 g, tempered samples (21°C), were monadically presented in black plastic cups with lids, labelled with randomized three-digit codes. Sensory responses were recorded on 15 cm anchored scales using FIZZ data acquisition software (Biosystems, Couternon, France). Samples were presented in a complete randomised block design over three replicated sessions. Still water at room temperature was provided for palate cleansing.

### 2.4. Chemical analysis

#### 2.4.1. Extraction of metabolites

Metabolites were extracted from snack bars and RPC by homogenizing 50 mg product by shaking with chrome balls at 30 hz in a Retch Mixer Mill for 2x5 minutes with 300 µL 85% methanol containing 5 µM p-hydroxybenzyl glucosinolate as internal standard. After homogenization, samples were incubated in a thermomixer compact from Eppendorf at 800 rpm for 60 min at room temperature (21°C). Subsequently, the samples were centrifuged at 10,000 g for 5 min and the supernatant was used for subsequent analysis.

#### 2.4.2. Metabolite LC-MS/MS analysis of rapeseed press cake

For the analysis of glucosinolates, flavonoids, phenylpropanoids and goitrin by liquid chromatography coupled to tandem mass spectrometry (LC-MS/MS), RPC extracts were diluted in deionized water 25- and 2500-fold, and energy bar extracts were diluted 5- and 500-fold. Four replicates for each sample were analysed, and analyte concentrations were determined by comparing their signal against calibration standard curves. Calibration curves were prepared for each analytical run using calibration standards prepared in deionized water at concentrations 1, 10, 50, 100, 200, 500, 1000, 2000, 5000 and 10000 nmol/L. p-hydroxybenzyl glucosinolate, which was added to the extraction solvent, was used as internal standard to control for analyte losses during extraction and inter-injection variation during analysis. Sinapine was detected in snack bars and RPC extract with a concentration above the limit of quantification. Further dilution of the RPC extract resulted in a quenching of the signal disproportionate to the dilution, suggesting that peak area is highly influenced by matrix effects in our experimental setup, thus making the measurements unreliable.

For free amino acid analysis by LC-MS/MS, extracts from both RPC and snack bars were diluted in deionized water 25-fold and subsequently 10-fold in 13C-, 15N-labelled Cell Free Amino Acid Mixture (Merck Sigma-Aldrich). Samples were run in four replicates, except for dehulled RPC for which only three replicates were used. Quantification of the individual free amino acids was performed from the 13C, 15N-labeled amino acids used as internal standards at known concentrations. See Supplementary Methods 1 and 2 for detailed descriptions of the LC-MS/MS methods.

#### 2.4.3. Quantification of crude protein

The total nitrogen content of raw RPC and snack bars were quantified by combustion in a vario MACRO cube CHNS analyzer from Elementar. To estimate the samples’ content of crude protein, a nitrogen-to-protein conversion factor of 6.25 was used (FAO/WHO, 1991).

### 2.5. Data analysis

The sensory profiling data were analysed with IBM SPSS by different ANOVA models. For evaluating individual sample differences in the GLM procedure a 2-way mixed-model ANOVA was used with the factors samples(fixed) and assessors(random) and their interaction. For analysing the effect of RPC dosage (7, 14, and 21%) and seed treatment (whole or dehulled) a 3-way mixed model ANOVA was used with the factors dosage (fixed), seed treatment (fixed) and assessor (random) and their 2-way interactions. In order to compare the control sample, a separate ANOVA was run excluding the factor seed treatment. In all ANOVA’s, Tukey’s test for comparisons of sample differences was used as post-hoc test with a significance level of α=0.05.

Principal Component Analysis (PCA) was conducted in R version 4.5.2 (R Core Team, 2025) with the package ‘FactoMineR’ (Lê et al., 2008). Mean values were used for each sample, and the data was standardized before analysis. The analysis used sensory attributes with significant differences as active variables and metabolites were added as supplementary variables. The results were visualised as a biplot applying the scaling of variables implemented in ’factoextra’ (Kassambara & Mundt, 2020). All visualisations were done using ‘tidyverse’ (Wickham et al., 2019).

## 3. Results

### 3.1 Sensory evaluation of RPC-containing snack bars

We formulated raw RPC-containing snack bars to investigate the sensory characteristics of a raw RPC-containing food product. The snack bars were based on whole food ingredients (including dates, raisins, nuts and coffee) with an increasing proportion (0%, 7%, 14%, and 21%) of raw RPC from whole or dehulled seeds (**Table 1**, **Figure 1**). A panel of trained tasters generated a vocabulary of thirteen sensory attributes that characterized the sensory impressions of raw RPC in the food product (**Table 2**). The panel subsequently rated the intensity of these sensory attributes in snack bars with different doses of RPC from whole or dehulled seeds (**Table 3**). The intensity of seven of eight dose-dependent attributes plateaued in snack bars with >14% RPC (**Figure 2)**. Intensities of oxidized odor and bitter taste were significantly higher in bars with 7% and 14% RPC than in the 0% control whereas cabbage odor, cabbage flavor, bitter aftertaste and astringent afterfeel did not increase at 7% RPC but first at 14% and 21%. Noticeably, the intensity of astringent mouthfeel was the only sensory attribute that increased proportionally with dose. With respect to impact of dehulling, bars containing RPC from dehulled seeds were rated higher for four sensory attributes (overall odor intensity, cabbage odor, cabbage flavor and popcorn flavor) compared with bars containing RPC from whole seeds (**Table 2**). Additionally, the intensity of the sensory attributes burned odor, sour taste and ryebread flavor as well as pungent mouthfeel detected in raw RPC, did not increase in snack bars in a dose- or dehulling-dependent manner, showing that RPC does not impact these sensory attributes to the food product significantly. Furthermore, the highest intensity scores (>10 out of 15) were recorded for overall intensity of odor, oxidized odor, bitter taste, bitter aftertaste and astringent mouthfeel. These results indicate that raw RPC markedly influenced the sensory profile of the snack bars in a dose-dependent manner, with the most pronounced changes occurring at <14% RPC, and that dehulling treatment has an impact on specific odor and flavor attributes in the snack bars.

**Figure 1:**
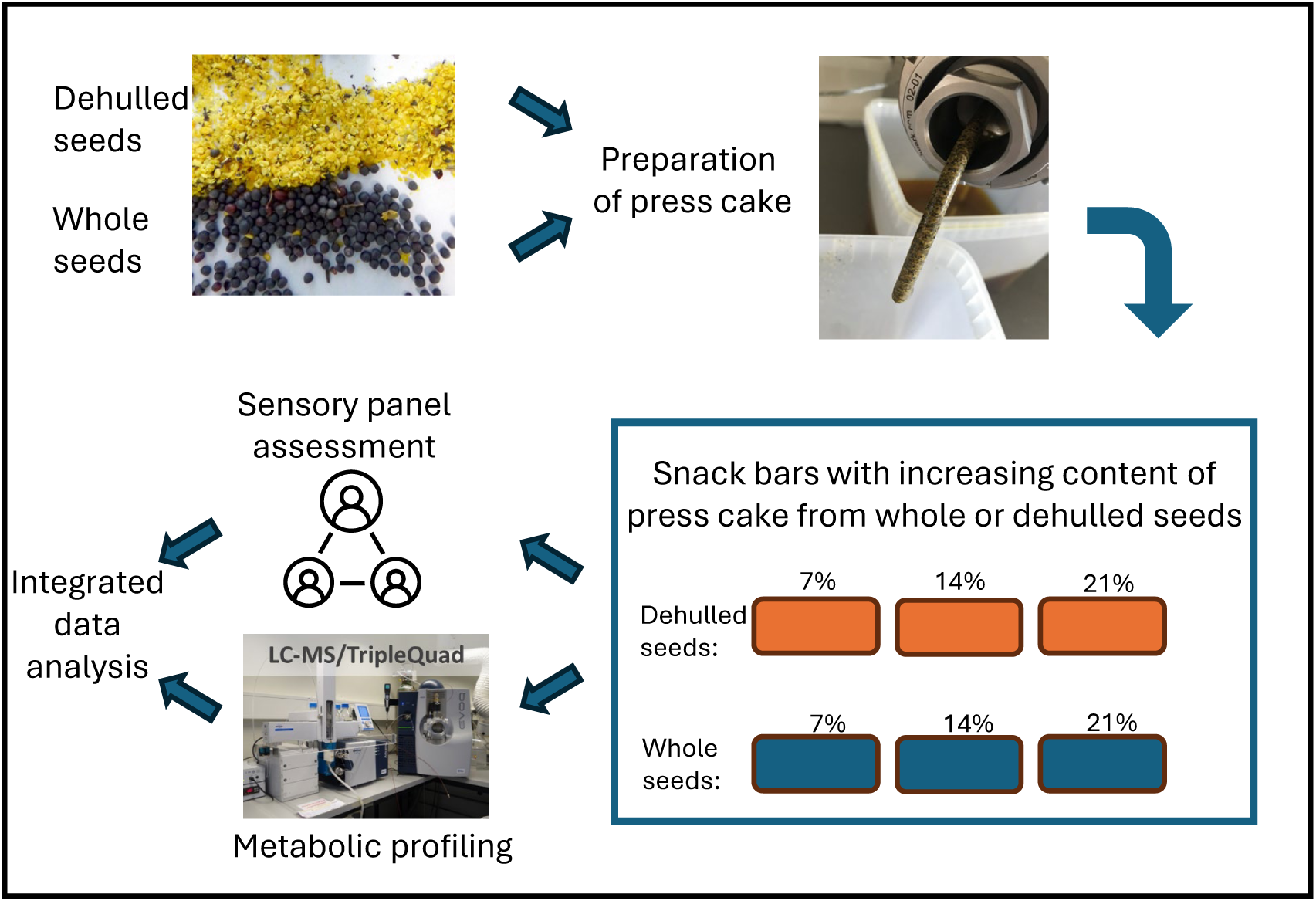
Workflow of study from preparation of press cake to integrated data analysis.

**Figure 2:**
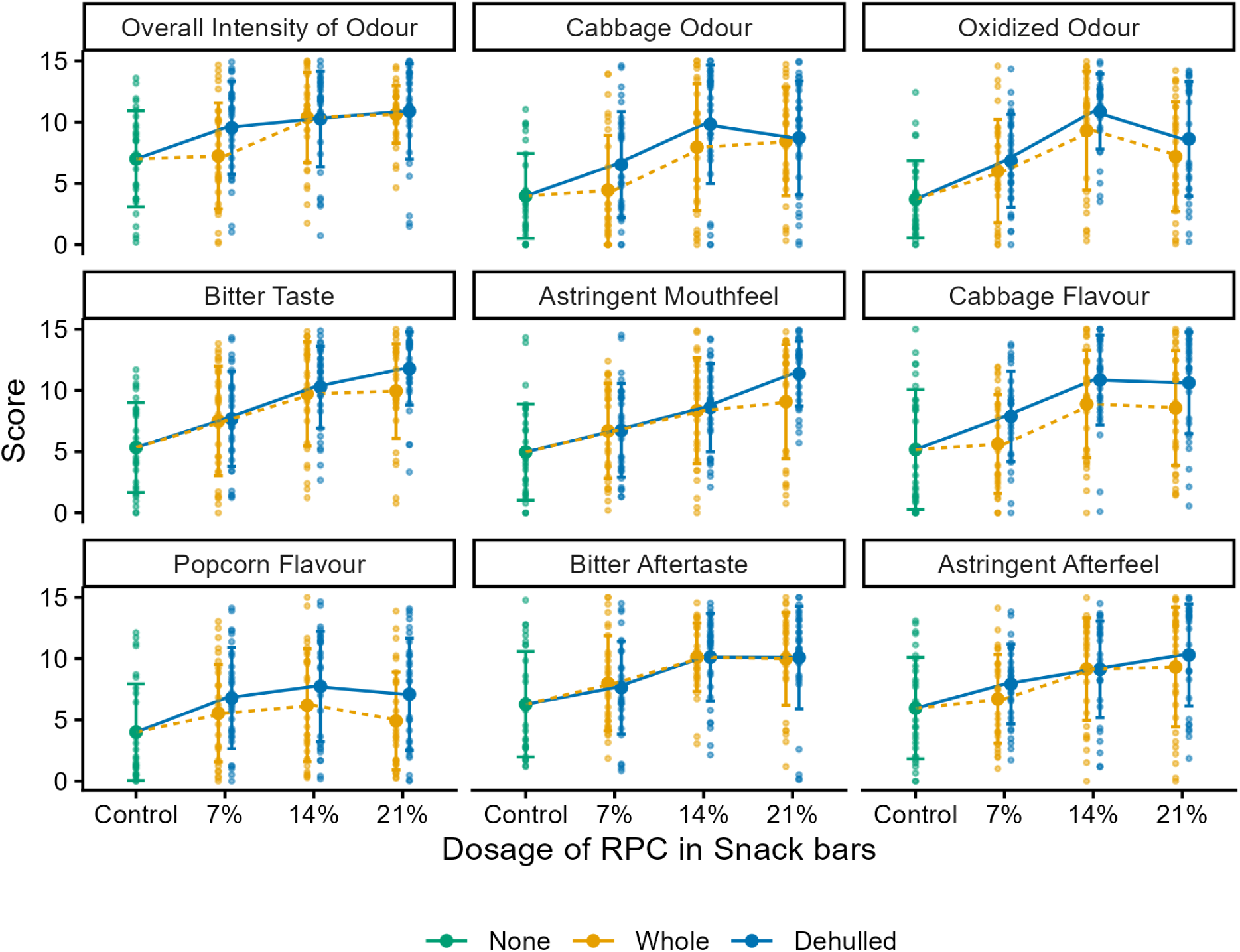
Intensity of sensory attributes in snack bars with press cake from whole dehulled seeds. For Overall intensity of Odour, the scale goes from 0-15 where 0 is eakest odour and 15 is stronger odour. For all other sensory attributes, 0 is a little and is a lot of the particular attribute. Small dots represent individual sensory easurements, large dots represent means of sensory scores, and error bars represent andard deviations.

**Table 2:** Vocabulary describing the sensory attributes characteristic of raw rapeseed press cake.

| Sensory attribute | Definition | Scale range | Reference material |
| --- | --- | --- | --- |
| **Odour** |  |  |  |
| Overall intensity | Strength of the sample's odor | Weak - strong | None |
| Burned | Odor of burnt or over-toasted food | A little – a lot | Burned toasted white sandwich bread |
| Cabbage | Odor reminiscent of raw, shredded white cabbage | A little – a lot | Shredded white cabbage |
| Oxidized | Odor associated with stale, rancid, or cardboard-like notes | A little – a lot | Aged almond oil |
| **Taste/flavor** |  |  |  |
| Sour | Taste associated with citric acid solution | A little – a lot | Citric acid (2 g/L water) |
| Bitter | Taste associated with quinine solution | A little – a lot | Quinine (1 g/L water) |
| Ryebread | Flavor characteristic of untoasted rye bread | A little – a lot | Untoasted ryebread slice |
| Cabbage | Flavor reminiscent of raw, shredded white cabbage | A little – a lot | Shredded white cabbage |
| Popcorn | Flavor reminiscent of freshly popped, unsalted popcorn | A little – a lot | Unsalted fresh popcorn |
| **Mouthfeel** |  |  |  |
| Astringent | Dry, puckering sensation on the tongue and inner cheeks and gums | A little – a lot | 2 black tea bags (Medova) infused in 250 mL water |
| Pungent | Irritating, sharp, mustard-like sensation in the nasal and oral cavity | A little – a lot | Fresh arugula leaves |
| **After impressions** |  |  |  |
| Bitter aftertaste | Lingering bitterness remaining after swallowing/ spitting, comparable to the aftertaste of black tea | A little – a lot | Quinine (1 g/L water) |
| Astringent afterfeel | Lingering dry, puckering sensation after swallowing/ spitting | A little – a lot | 2 black tea bags (Medova) infused in 250 mL water |

**Table 3:** Statistical analysis of whether dose and dehulling treatment affect sensory attributes. Sensory panel mean intensity values for snack bars containing different dosages of rapeseed press cake (RPC) made from, respectively, whole or dehulled seeds and attributes showing product differences. Abbreviations: ‘Whole’ snack bars with RPC from whole seeds; ‘Dehulled’: snack bars with RPC from dehulled seeds.

| Sensory attribute | Dosage |  |  |  | Dehulling Treatment |  | Dosage |  | Dehulling Treatment |  | Dosage x Dehul. Tr. |  |
| --- | --- | --- | --- | --- | --- | --- | --- | --- | --- | --- | --- | --- |
|  | 0%<sup>a</sup> | 7% | 14% | 21% | Whole | Dehulled | F(2,18) | *p*-value | F(1,9) | *p*-value | F(2,138) | *p*-value |
| **Odor** |  |  |  |  |  |  |  |  |  |  |  |  |
| Overall intensity | **7.0b** | **8.4b** | **10.3a** | **10.8a** | **9.4b** | **10.2a** | 6.570 | **0.007** | 8.821 | **0.016** | 2.518 | 0.084 |
| Burned | 8.1 | 8.3 | 8.2 | 8.6 | 8.7 | 8.0 | 0.140 | 0.870 | 0.744 | 0.411 | 1.342 | 0.265 |
| Cabbage | **4.0b** | **5.5b** | **8.9a** | **8.6a** | **7.0b** | **8.4a** | 8.770 | **0.002** | 9.038 | **0.015** | 0.828 | 0.439 |
| Oxidized | **3.7c** | **6.4b** | **10.1a** | **7.9b** | 7.5 | 8.8 | 6.316 | **0.008** | 3.613 | 0.090 | 0.165 | 0.848 |
| **Taste/Flavor** | 7.4 | 7.7 | 7.9 | 7.7 | 7.2 | 8.3 | 0.062 | 0.940 | 4.998 | 0.052 | 2.333 | 0.101 |
| Sour |  |  |  |  |  |  |  |  |  |  |  |  |
| Bitter | **5.3c** | **7.6b** | **10.0a** | **10.9a** | 9.1 | 9.9 | 10.937 | **<0.001** | 1.408 | 0.266 | 1.065 | 0.347 |
| Ryebread | 8.0 | 8.6 | 7.9 | 7.8 | 8.4 | 7.8 | 1.074 | 0.363 | 1.258 | 0.291 | 0.064 | 9.938 |
| Cabbage | **5.2b** | **6.8b** | **9.9a** | **9.6a** | **7.7b** | **9.8a** | 8.034 | **0.003** | 36.510 | **<0.001** | 0.028 | 0.973 |
| Popcorn | **4.0b** | **6.2a** | **7.0a** | **6.0a** | **5.6b** | **7.2a** | 0.933 | 0.411 | 5.434 | **0.045** | 0.266 | 0.766 |
| **Mouthfeel** |  |  |  |  |  |  |  |  |  |  |  |  |
| Astringent | **5.0d** | **6.7c** | **8.5b** | **10.2a** | 8.0 | 8.9 | 7.692 | **0.004** | 2.013 | 0.190 | 2.480 | 0.087 |
| Pungent | 5.5 | 6.2 | 6.5 | 6.5 | 6.3 | 6.5 | 0.072 | 0.931 | 0.089 | 0.773 | 0.489 | 0.615 |
| **After impressions** |  |  |  |  |  |  |  |  |  |  |  |  |
| Bitter aftertaste | **6.3b** | **7.8b** | **10.1a** | **10.0a** | 9.4 | 9.3 | 3.806 | **0.042** | 0.038 | 0.850 | 0.103 | 0.902 |
| Astringent afterfeel | **6.0b** | **7.3b** | **9.1a** | **9.8a** | 8.4 | 9.1 | 3.808 | **0.042** | 1.304 | 0.283 | 0.837 | 0.435 |

#### 3.2.1. Metabolites associated with bitterness in raw RPC

Next, we generated a list of known bitter compounds that are found in seeds of rapeseed (phenolics, flavonoids, glucosinolates and their hydrolysis products) (**Supplementary Table 1**). In the raw RPC, sinapic acid concentration was approximately 7.5–10 µmol/g, with whole seed RPC generally at the higher end of this range. In the snack bars, sinapic acid levels increased with proportion of RPC (**Figure 3A**). Bars made with 21% RPC from dehulled seeds contained sinapic acid in proportion to the amount of RPC added, whereas all other tested snack bars contained less-than-expected sinapic acid (**Figure 3A**). Unfortunately, attempts to quantify sinapine in the samples were unsuccessful most likely due to severe matrix effects.

**Figure 3:**
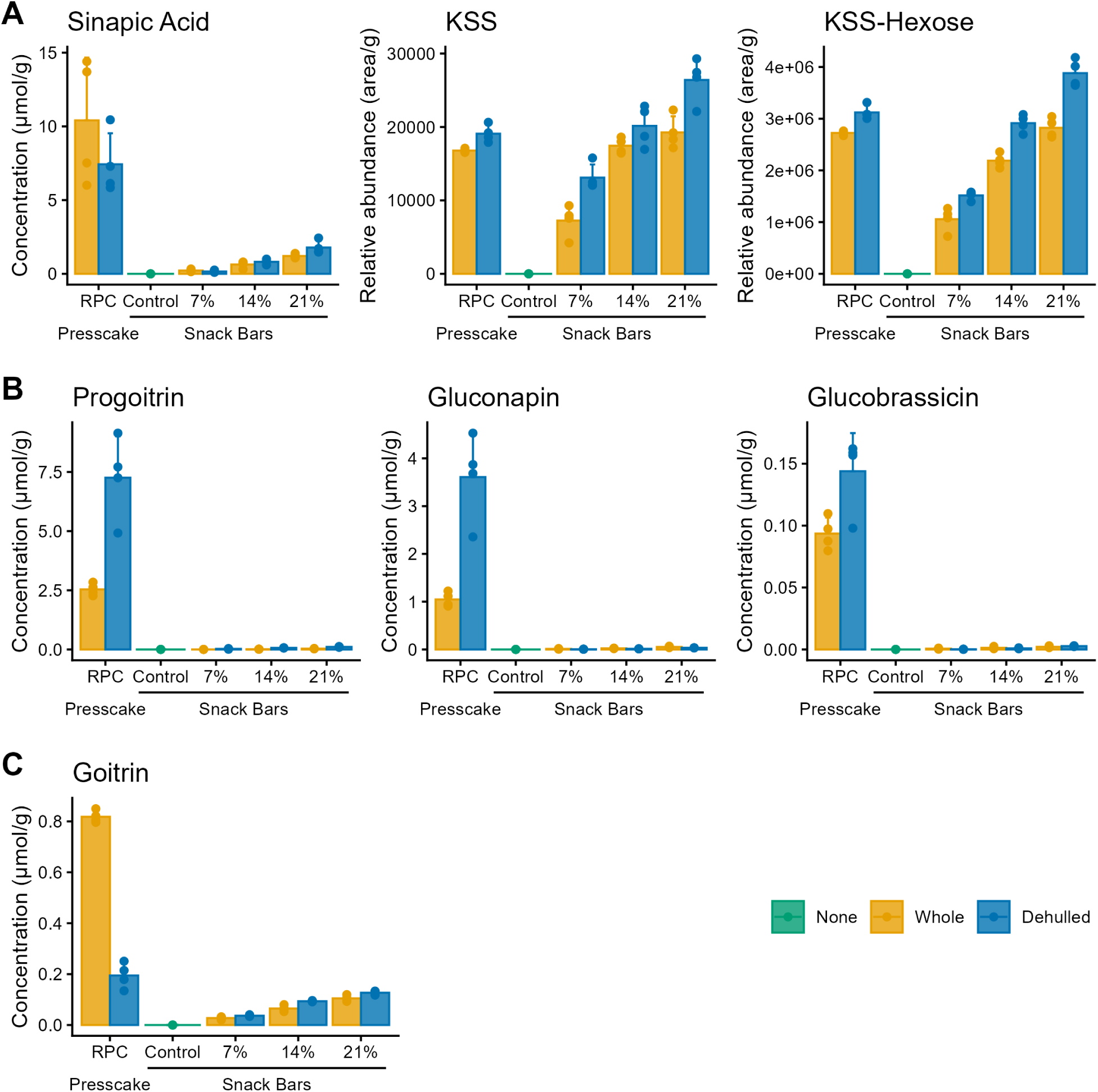
Quantification of known bitter compounds in raw rapeseed press cake (RPC) and snack bars supplemented with RPC. Note that the values for KSS and KSS-Hexose are peak areas normalized to sample weight. The percentages refer to the dosage of RPC included in the snack bars (mass percentage). The top of the bars are averages of four technical replicates and error bars show the standard deviation for the technical replicates. KSS: kaempferol-3-o-sinapoylsophoroside, KSS-Hexose: kaempferol-3-o-sinapoylsophorosideglucoside.

We quantified the key bitter compounds KSS and a KSS-hexose, known from rapeseed protein isolates, to investigate how introducing RPC into snack bars affected the profile of these compounds. We found that KSS and KSS-hexose were present in both raw RPC and snack bars and that peak area of both compounds increased with increasing dosage of RPC in snack bars (**Figure 3A**). No KSS or KSS-hexose was detected in control snack bars without RPC. Surprisingly, the peak area of KSS and KSS-hexose in snack bars was much higher than expected based on the peak areas observed in analysis of raw RPC (respectively, 5-9-fold or 5-7-fold higher). This suggests that KSS and KSS-hexose may be generated from higher order derivatives during preparation of the snack bar or be more easily extractable from the snack bar food matrix.

We quantified three glucosinolates (progoitrin, gluconapin, glucobrassicin) selected for their abundance and documented bitter taste as well as one hydrolysis product, goitrin (derived from progoitrin) also with reported bitter taste (**Supplemental Table 1**). In raw RPC from whole seeds, the sum of quantified glucosinolates was 4.08 µmol/g, and of these progoitrin was the most abundant glucosinolate with a concentration of 2.54 µmol/g, followed by gluconapin at 1.05 µmol/g and glucobrassicin at 0.49 µmol/g, respectively (**Figure 3B**). The level of goitrin was 0.82 µmol/g (**Figure 3C**). In the raw RPC from dehulled seeds, the total level of quantified glucosinolates was 12.74 µmol/g, i.e. ∼3 fold higher than RPC from whole seeds, with progoitrin being most abundant with a concentration of 7.26 µmol/g, followed by gluconapin at 3.61 µmol/g and glucobrassicin at 1.87 µmol/g, respectively (**Figure 3B**). The level of goitrin was 0.195, i.e. ∼4x lower than the level found in RPC from whole seeds (**Figure 3C)**. This shows that RPC from whole seeds have less of the selected glucosinolates than RPC from dehulled seeds but show signs of significant glucosinolate hydrolysis taking place prior to incorporation into the food matrix.

In the snack bars with 7%, 14%, or 21% raw RPC from whole seeds, we observed progoitrin levels of 0.046, 0.057, and 0.137 µmol/g, respectively, whereas the snack bars with RPC from dehulled seeds contained 0.023, 0.064 and 0.101 µmol/g, respectively (**Figure 3B**). These results show that upon preparation of the snack bars and thereby incorporation of RPC into a food matrix, extensive hydrolysis of glucosinolates occurs. Unexpectedly, only small amounts of goitrin were detected in the snack bars, accounting for 7.3-15.6% of the progoitrin added to the snack bars (**Figure 3C**).

### 3.3 Correlation between sensory scores and content of RPC-derived metabolites

Next, we investigated whether the variation in intensities of sensory attributes between samples correlated with the content of RPC-derived bitter compounds in the snack bars. Based on a principal component analysis, we generated a biplot combining dose- and dehulling-dependent sensory attributes with measured metabolites added as supplementary variables. The PCA biplot indicated that the snack bars formed two distinct clusters along PC1 (86.7%): 0–7% RPC and 14–21% RPC, based on correlations among dose- or dehulling treatment-dependent sensory attributes and RPC content (Figure 4). All attributes were observed to be driving this separation, except the popcorn flavor. The overall odor intensity, bitter taste, bitter aftertaste, astringent mouthfeel and astringent afterfeel was positively associated with content of KSS, KSS-hexose and goitrin in the snack bars (**Figure 4**). The content of sinapic acid, gluconapin and glucobrassicin showed weaker association with any dose- or dehulling-dependent sensory attributes than the bitter compounds goitrin, KSS, and KSS-hexose, suggesting that in the food matrix these metabolites are not key for the intensity of sensory attributes characteristic of RPC. The cabbage odor was associated with the content of progoitrin in snack bars. Over PC2 (7.52%), we observed a trend of separation between RPC from whole or dehulled seeds. When comparing snack bars with the same dose of RPC from whole or dehulled seeds, bars containing press cake from dehulled seeds were more strongly associated with popcorn flavor and, to a lesser extent, oxidized odor. In conclusion, variation in the sensory profile of the snack bars was primarily associated with RPC concentration and the levels of goitrin, KSS, and KSS-hexose, while the modest dehulling-related separation observed along PC2 was not associated with the metabolites quantified in this study.

**Figure 4:**
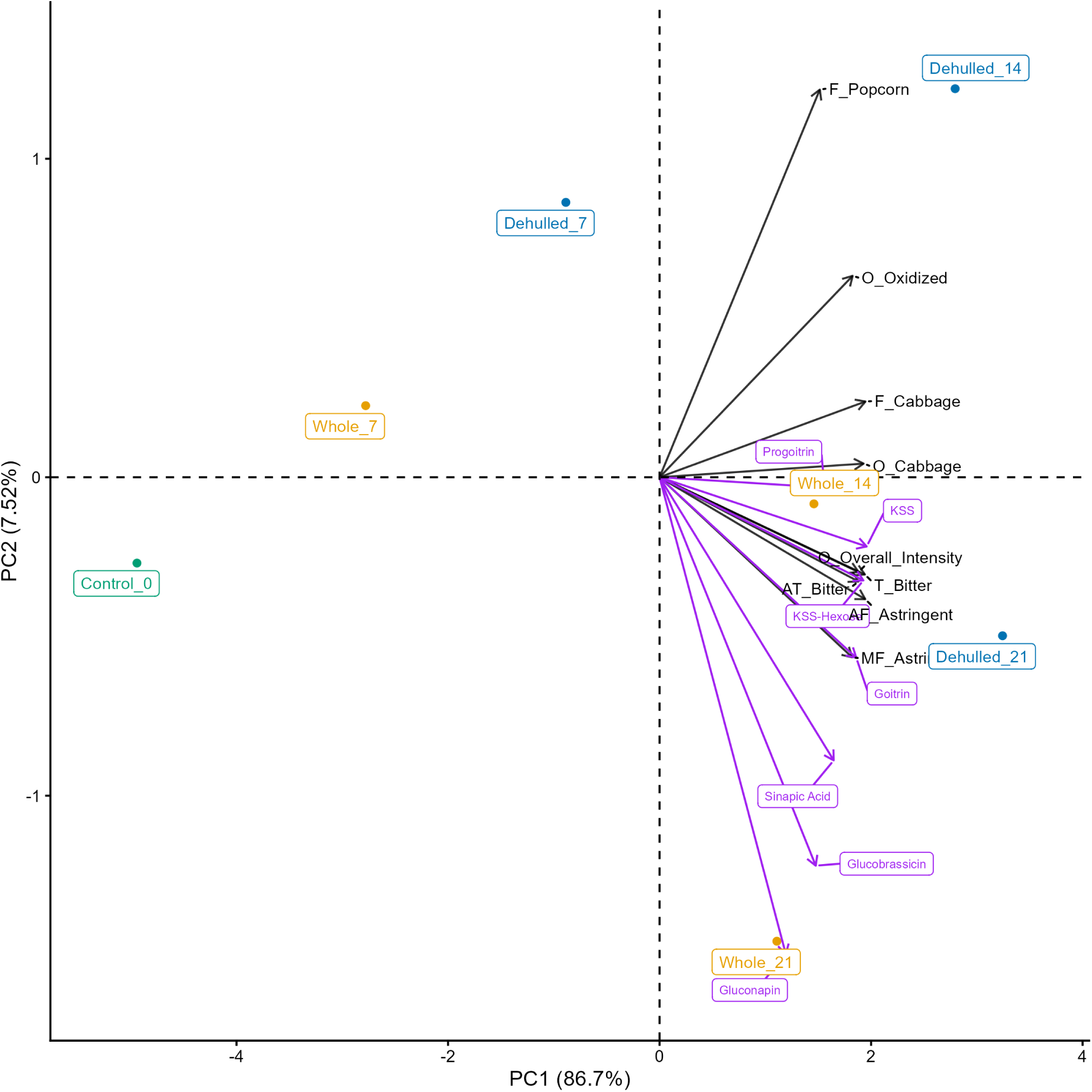
Biplot of sensory attributes with bitter metabolites projected as supplementary variables. Snack bars corresponding to sample IDs are detailed in **Table 1**. Abbreviations: O: Odour, T: Taste, F: Flavour, MF: Mouthfeel, AT: Aftertaste, AF: After Mouthfeel.

### 3.4. Dose-over-thresholds values

A bitterness threshold - the concentration at which half of a tasting panel can perceive a compound’s bitterness - can be used to calculate the DoseoverThreshold (DoT) factor, which indicates whether the compound is present at levels high enough to influence a food’s flavour. For progoitrin, the DoT factors for all snack bars were <0.1, while for goitrin, the DoT factors were in the range 0.3-1.4 (dependent on dose and dehulling treatment, **Table 4**). This observation suggests that goitrin, but not progoitrin, contribute to RPC-derived bitterness in the snack bars.

**Table 4:**
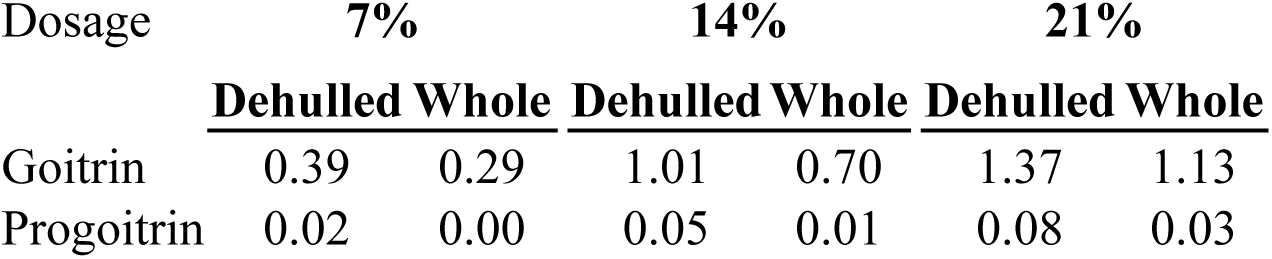
Dose-over-Threshold factors (DoT-factors) for goitrin and progoitrin in snack bars containing rapeseed press cake (RPC). A DoT-Factor of more or equal to 1 suggests that the particular compound should be detectable as bitter based on the perception in an aqueous solution. The bitterness threshold values are extracted from observations by Fenwick et al. (1983). ‘Dehulled’ and ‘Whole’ refers to the source of RPC used in the formulation of the particular snack bar.

### 3.5. Protein and amino acids

We analyzed crude protein and free amino acids in the RPC and snack bars. Raw RPC from whole and dehulled seeds contained 28% and 32% crude protein, respectively (Supplementary Figure 1). The free amino acid profile of both materials was dominated by glutamate (3-3.5 µmol/g), followed by alanine (∼1.0 µmol/g) and aspartate (0.5-0.8 µmol/g) (Supplementary Figure 2). Concentrations of individual free amino acids were generally slightly higher in RPC from dehulled than from whole seeds.

Incorporation of RPC into snack bars at 7-21% did not substantially affect total crude protein content, which remained at approximately 15-17% across all formulations (Supplementary **Figure 1**). The free amino acid composition of the snack bars differed markedly from that of the raw RPC. In all snack bar samples, proline was the predominant free amino acid (3.3-6.6 µmol/g), followed by alanine (1.8-2.3 µmol/g) and glutamine (0.7-0.9 µmol/g), whereas the remaining amino acids were present at lower concentrations. Although incorporation of RPC resulted in variation in concentration of several individual amino acids, no clear dose or dehulling treatment-dependent effect was observed. These show that inclusion of RPC in snack bars did not alter the crude protein content of the snack bars and only had a limited effect on the profile of free amino acids.

## 4. Discussion

The present work adopts a novel approach to characterize the sensory contribution of raw RPC in a food matrix and - by combining sensory and targeted metabolomic data - generates new insights on how bitter rapeseed-derived metabolites contribute to sensory attributes. The sensory contribution of RPC was dominated by oxidized and cabbage odors, bitter taste and cabbage flavor as well as astringent mouthfeel and bitter aftertaste. The intensity of most RPC-derived attributes was dose-dependent, with astringent mouthfeel notably increasing its intensity almost linearly with dose. For other dose-dependent attributes, addition of more than 14% RPC to the snack bars resulted in only a minimal increase in perceived intensity. The dose-response plateaus may reflect sensory adaptation. The non-linear effects may arise from matrix interactions, by which some bitter compounds may be partially masked as their concentration rises in the dense and sweet snack bar food matrix. Notably, retainment of the seed hull (i.e. using RPC from whole seeds) did not increase bitterness. This indicates that metabolites enriched primarily in the embryo are responsible for RPC-derived sensory attributes detectable in the snack bar food matrix. In fact, bars with RPC from dehulled seeds were perceived to be more intense for some attributes. A possible explanation is that hull fibres bind metabolites responsible for sensory attributes. Alternatively, the hulls themselves may serve as a dilution factor of RPC, reducing effective concentration. Furthermore, the concentration of protein was not notably higher in snack bars with RPC from dehulled seeds versus whole seeds. Together, the results indicate that there is no apparent advantage in terms of sensory impact of RPC or content of crude protein in dehulling seeds prior to oil pressing.

The measurements of selected known bitter compounds show a tight correlation of intensity of bitter taste and the peak area of KSS, KSS-hexose and the concentration of sinapic acid and goitrin (**Figure 4**), suggesting that these metabolites contribute to perceived bitterness of raw RPC in the snack bar food matrix. The observation that certain snack bars had a DoT-factor values above 1 for goitrin provides the first evidence that goitrin may contribute to RPC-derived bitterness in an RPC-containing food matrix. This finding is consistent with earlier observations reported for goitrin in Brussels sprouts (Fenwick et al., 1983). At highest RPC levels (21%) intensity of bitter taste and bitter aftertaste did not rise proportionally with the increased concentration of these metabolites, presumably due to masking of the additional bitterness by the complex food matrix. Our study provides evidence that other compounds beside previously the reported sinapine and choline chloride (Ismail et al., 1981) may contribute to perceived bitterness of raw RPC.

For the selected glucosinolates and sinapic acid, the measured concentrations in the snack bars were lower than expected if dilution were the only factor. The observation that only ∼10% of the RPC-derived progoitrin was recovered as goitrin is noticeable. Goitrin is key for the anti-thyroid effect of RPC (Galanty et al., 2024) and unlike inorganic thiocyanate ions - a hydrolysis product of several glucosinolates - the goitrogenic effect is not alleviated by adequate iodine status (Thorsen et al., 2025). Possible explanations could be that goitrin has been further metabolized/derivatized or that goitrin is not efficiently extracted from the matrix with the employed protocol. The finding that the peak areas of KSS and KSS-hexose in the bars were far higher than expected from raw RPC levels (5-9x) suggests that higher order kaempferol conjugates were hydrolyzed into KSS or KSS-hexose during preparation of the food, or that the metabolites were more efficiently extracted from the snack bar food matrix compared to the raw RPC. Our targeted metabolite analysis indicates that goitrin, sinapic acid and kaempferol glycosides in rapeseed is at least partly responsible for the RPC-derived bitterness, astringency and overall odor intensity of up to mid-range doses. The lack of correlations between the dose-dependent sensory attributes like cabbage and popcorn flavor as well as oxidized odor of RPC and the selected bitter compounds, highlights a need for further studies using untargeted metabolomics.

Our study illustrates the need for chemical characterization of the final food product itself, since metabolites in the raw RPC undergo chemical changes when added to the food matrix. Furthermore, the results may be used by food scientists and chefs as inspiration to introduce RPC in their recipes and by breeders looking to breed new rapeseed varieties with improved sensory characteristics e.g. by breeding for reduced level of bitter compounds in the seed.

## ACKNOWLDGEMENTS

We would like to thank Louise Svenningsen for valuable technical assistance, Juan Agustin Muñoz from ‘Spora’ for development of snack bar recipe, Jan Günther for purifying KSS and the KSS-hexose derivate from rapeseed press cake and ‘Bornholms Oliemølle A/S’ for providing whole and dehulled rapeseed seeds.

## AUTHOR CONTRIBUTIONS

**Jakob Skytte Thorsen:** Methodology, Formal analysis, Investigation, Resources, Writing original draft, review and editing, project administration.

**Arendse Maria Toft:** Conceptualization, Funding acquisition, Formal analysis, Visualization, Writing.

**Andrea Bononad-Olmo:** Methodology, Formal analysis, Investigation, Writing – original, review & editing.

**Niels Christian Holm Sanden:** Conceptualization, Methodology, Project administration, Funding Acquisition, writing – review & editing.

**Kwadwo Gyapong Agyenim-Boateng:** Methodology, Data Curation.

**Michal Poborsky:** Investigation, Methodology, Data Curation.

**Christoph Crocoll:** Methodology,

**Barbara Ann Halkier:** Writing - review and editing, Conceptualization, Formal Analysis, Supervision, funding acquisition, Visualization.

**Wender L.P. Bredie:** Writing, Conceptualization, Formal Analysis, Supervision,

**Deyang Xu:** Conceptualization, Methodology, writing, supervision, Funding acquisition.

## Supplementary Methods

### Supplementary Method 1: Detection and quantification of glucosinolates, flavonoids and goitrin by LC-MS/MS

The method was modified from Crocoll et al. (2016) to include non-glucosinolate metabolites as well as to adjust and optimize parameters for the LC-MS/MS system in use. Briefly, chromatography was performed on a 1290 Infinity II UHPLC system (Agilent Technologies). Separation was achieved on a Kinetex XB-C18 column (100 x 2.1 mm, 1.7 µm, 100 Å, Phenomenex, Torrance, CA, USA). Formic acid (0.05%, v/v) in water and acetonitrile (supplied with 0.05% formic acid, v/v) were employed as mobile phases A and B, respectively. The elution profile was: 0-0.5 min, 3% B; 0.5-4.0 min, 3-45% B; 4.0-4.2 min 45-100% B, 4.2-5.1 min 100% B, 5.1-5.2 min, 100-3% B and 5.2-6.2 min 3% B. The mobile phase flow rate was 400 µL/min. The column temperature was maintained at 40°C. The liquid chromatography was coupled with Ultivo triple quadrupole mass spectrometer (Agilent Technologies) equipped with a Jetstream electrospray ion source (ESI). The instrument parameters were optimized by infusion experiments with pure standards, when available. The ion spray voltage was set to 4000V in positive mode and 3500V in negative mode. Dry gas temperature was set to 325°C and dry gas flow to 12 L/min. Sheath gas temperature was set to 400 °C and sheath gas flow to 12 L/min. Nebulizing gas was set to 50 psi. Nitrogen was used as dry gas, nebulizing gas and collision gas. Multiple reaction monitoring (MRM) was used to monitor precursor ion → fragment ion transitions. MRM transitions were determined by direct infusion experiments of reference standards except for kaempferol-3-O-sinapoyl-sophoroside (KSS) and kaempferol-3-O-sinapoylsophoroside-hexose (KSS-Hexose), where purified putative compounds were used, isolated from RPC by preparative LC. Both Q1 and Q3 quadrupoles were maintained at unit resolution. MRM transitions, fragmentor voltage and collision energies are detailed in Supplementary Table 2. Mass Hunter Quantitation Analysis for QQQ software (Version 10, Agilent Technologies) was used for data processing.

### Supplementary Method 2: Detection and quantification of free amino acids by LC-MS/MS

The same LC-MS system equipped with Zorbax Eclipse XDB-C18 column (150 x 3.0 mm, 1.8 µm, Agilent Technologies) was used for amino acid analysis. Formic acid (0.05 %, v/v) in water and acetonitrile (supplied with 0.05 % formic acid, v/v) were employed as mobile phases A and B, respectively. The elution profile was: 0.00-0.75 min, 3 % B; 0.75-4.10 min, 3-65 % B; 4.10-4.20 min, 65-100 % B; 4.20-4.95 min, 98 % B; 4.95-5.00 min, 100-3 % B; and 5.0-6.0 min 3 % B. The mobile phase flow rate was 400 µL/min. The MS instrument parameters were optimized by infusion experiments with pure standards. The ion spray voltage was set to 2500 V in a positive ion mode. Dry gas temperature was set to 325 °C and dry gas flow to 11 L/min. Sheath gas temperature was set to 350 °C and sheath gas flow to 12 L/min. Nebulizing gas was set to 40 psi. Nitrogen was used as dry gas, nebulizing gas and collision gas. MRM was used to monitor precursor ion → fragment ion transitions. MRM transitions and other parameters such as fragmentor voltage and collision energies were optimized using reference standards. Both Q1 and Q3 quadrupoles were maintained at unit resolution. MRM transitions, fragmentor voltage and collision energies are detailed in Supplementary Table 3. Mass Hunter Quantitation Analysis for QQQ software (Version 10, Agilent Technologies) was used for data processing See supplementary table 1 for an overview of the quantified metabolites.

**Supplementary Figure 1:**
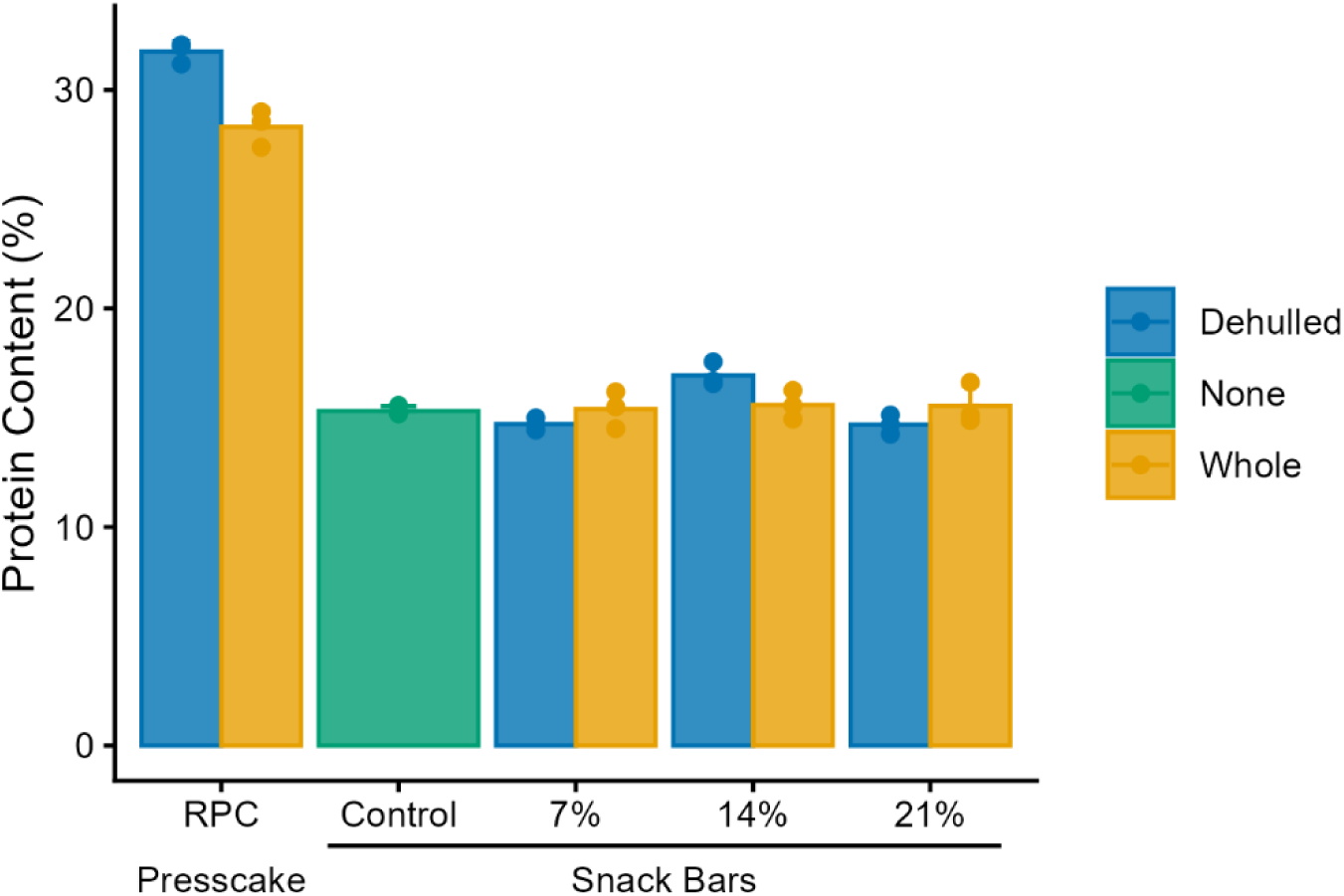
Concentration of crude protein in raw rapeseed press cake and snack bars. The top of the bars are averages of the technical replicates and error bars show the standard deviation for the technical replicates.

**Supplementary Figure 2.**
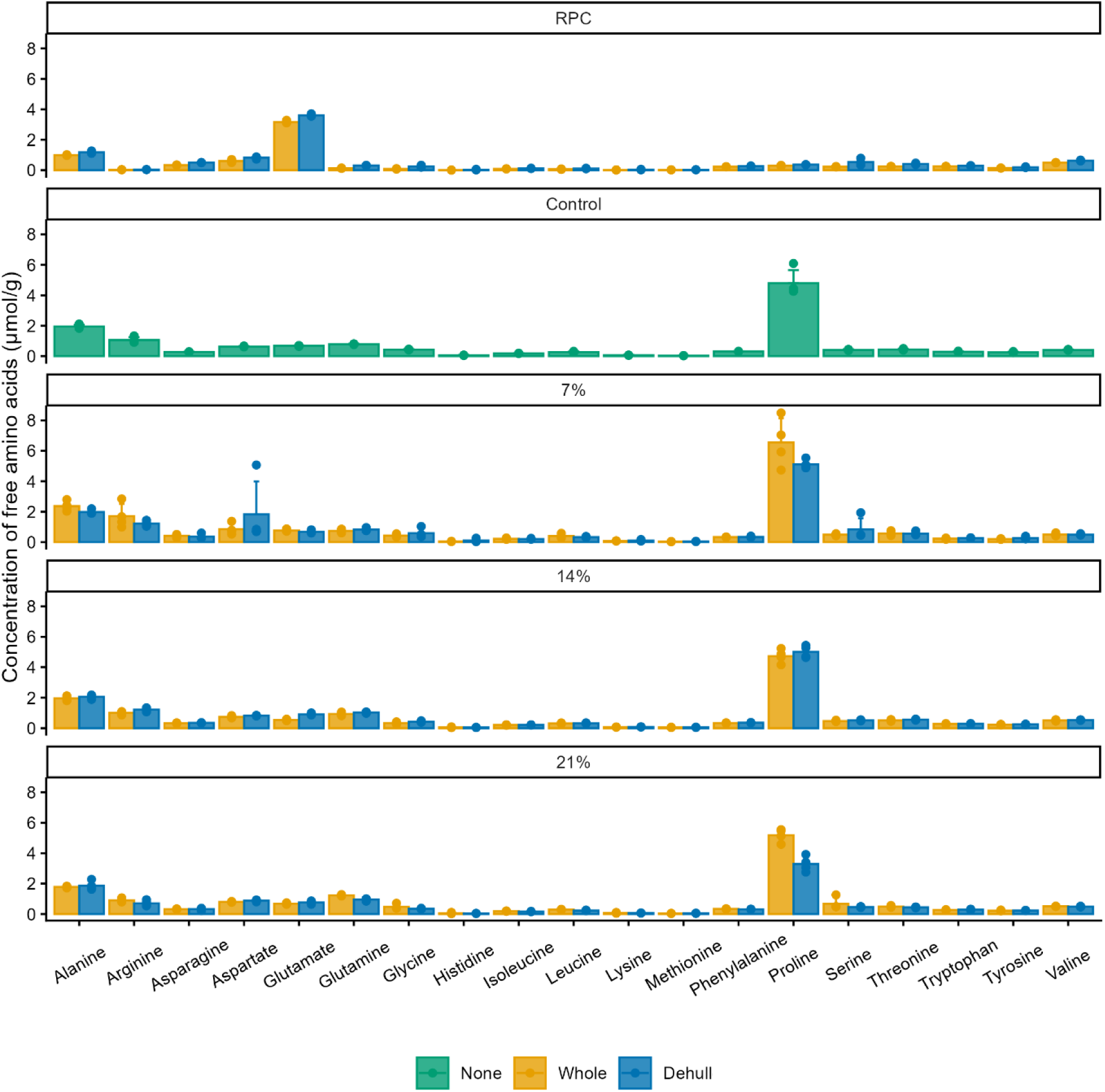
Free amino acids in raw rapeseed press cake and snack bars containing varying doses of RPC from ‘whole’ or ‘dehulled’ seeds. The top of the bars are averages of the technical replicates and error bars show the standard deviation for the technical replicates.

**Supplementary Table 1.**
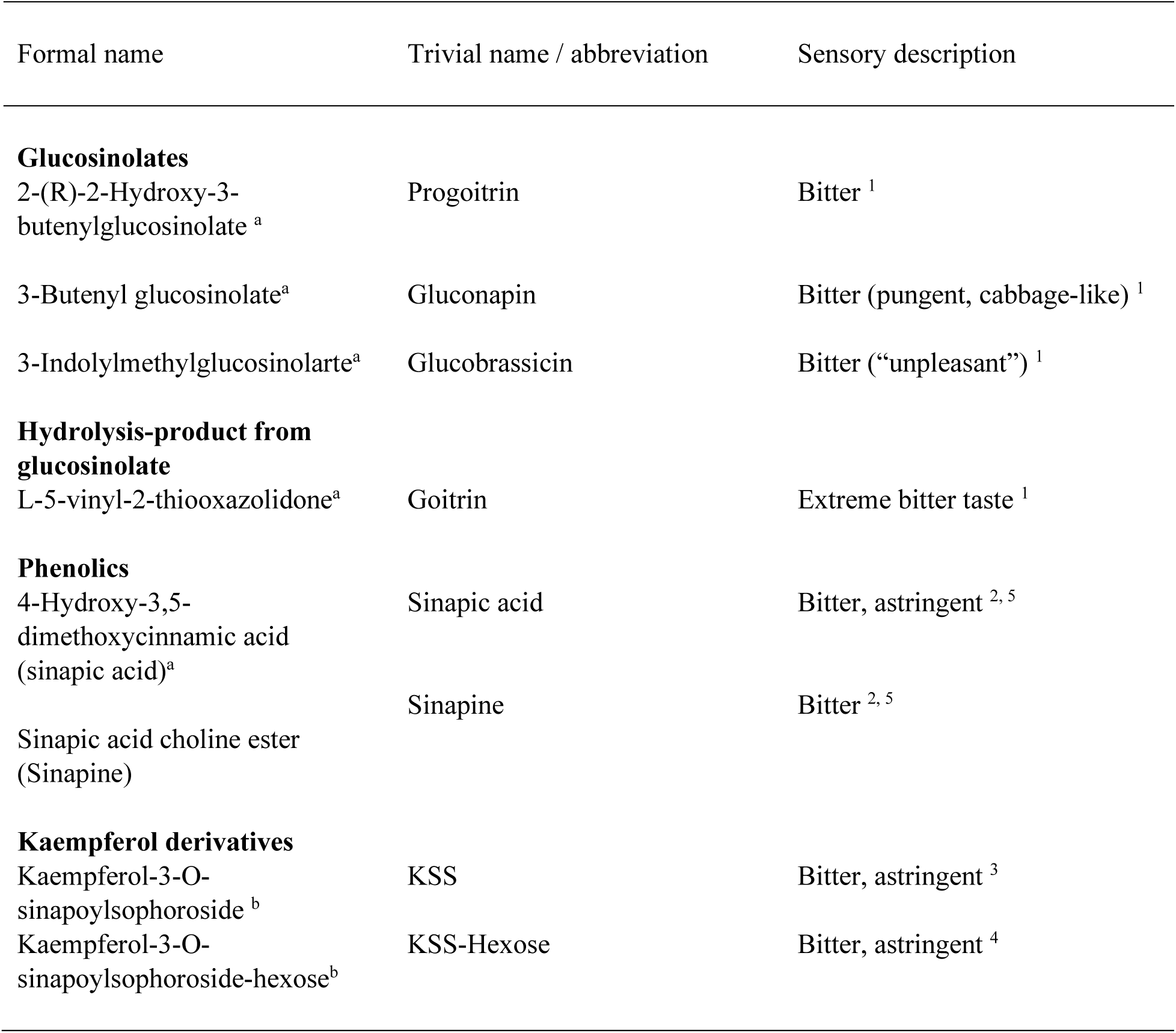
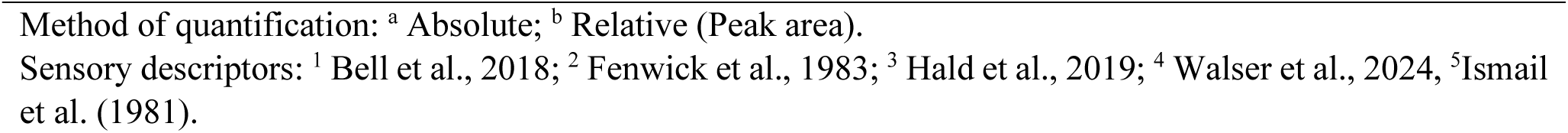
Overview of specialised metabolites identified using LC-MS.

| Formal name | Trivial name / abbreviation | Sensory description |
| --- | --- | --- |
| **Glucosinolates** |  |  |
| 2-(R)-2-Hydroxy-3-butenylglucosinolate<sup>a</sup> | Progoitrin | Bitter<sup>1</sup> |
| 3-Butenyl glucosinolate<sup>a</sup> | Gluconapin | Bitter (pungent, cabbage-like)<sup>1</sup> |
| 3-Indolylmethylglucosinolarte<sup>a</sup> | Glucobrassicin | Bitter (“unpleasant”)<sup>1</sup> |
| **Hydrolysis-product from glucosinolate** |  |  |
| L-5-vinyl-2-thiooxazolidone<sup>a</sup> | Goitrin | Extreme bitter taste<sup>1</sup> |
| **Phenolics** |  |  |
| 4-Hydroxy-3,5-dimethoxycinnamic acid (sinapic acid)<sup>a</sup> | Sinapic acid | Bitter, astringent<sup>2, 5</sup> |
| Sinapic acid choline ester (Sinapine) | Sinapine | Bitter<sup>2, 5</sup> |
| **Kaempferol derivatives** |  |  |
| Kaempferol-3-O-sinapoylsophoroside<sup>b</sup> | KSS | Bitter, astringent<sup>3</sup> |
| Kaempferol-3-O-sinapoylsophoroside-hexose<sup>b</sup> | KSS-Hexose | Bitter, astringent<sup>4</sup> |

Method of quantification: <sup>a</sup> Absolute; <sup>b</sup> Relative (Peak area).
Sensory descriptors: <sup>1</sup> Bell et al., 2018; <sup>2</sup> Fenwick et al., 1983; <sup>3</sup> Hald et al., 2019; <sup>4</sup> Walser et al., 2024, <sup>5</sup> Ismail et al. (1981).

**Supplementary Table 2:**
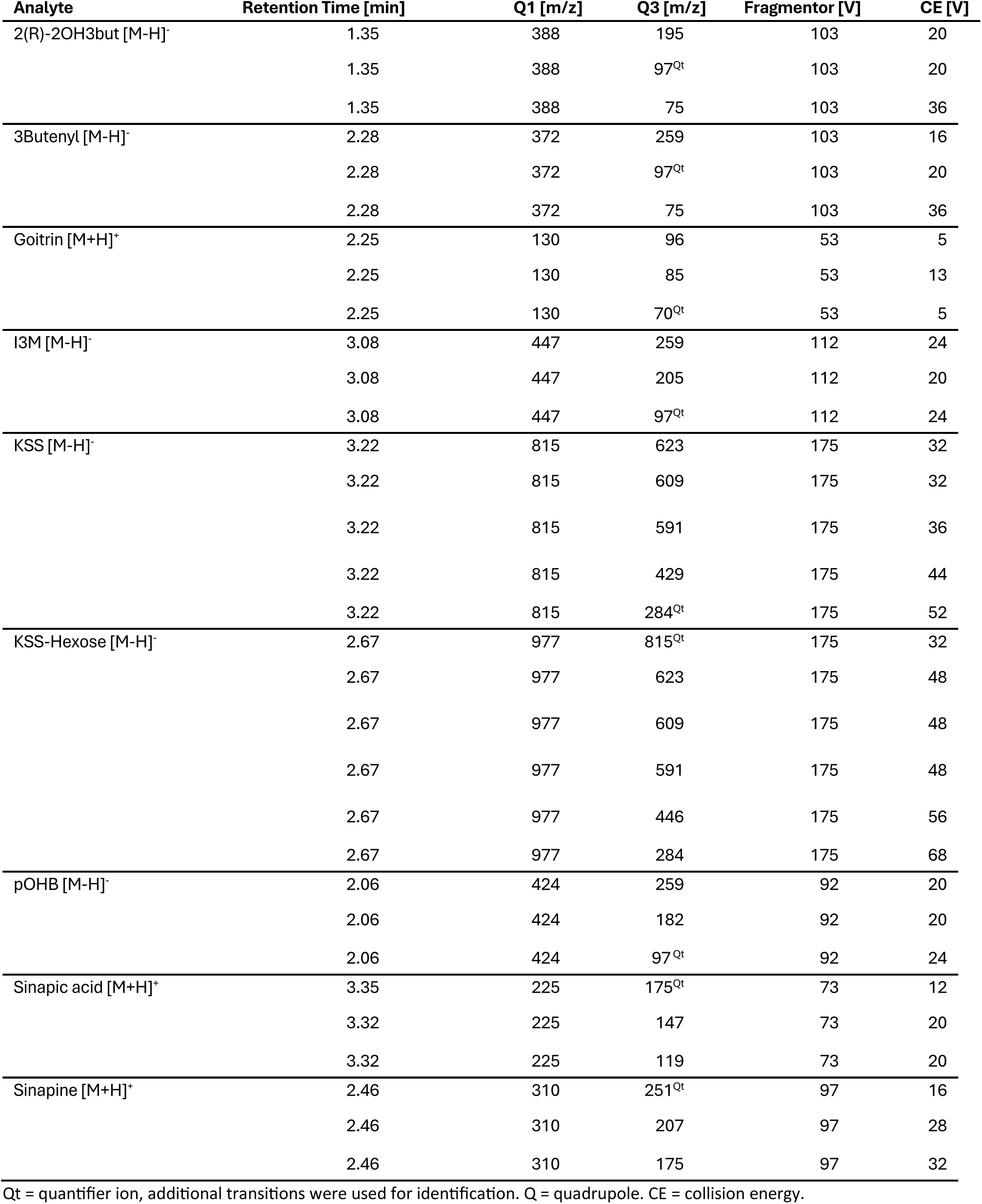
MRM transitions, fragmentor voltages and collision energies used for LC-MS/MS.

**Supplementary Table 3:** MRM transitions, fragmentor voltages and collision energies used for amino acid LC-MS/MS analysis.

| Analyte | Retention Time [min] | Q1 [m/z] | Q3 [m/z] | Fragmentor [V] | CE [V] |
| --- | --- | --- | --- | --- | --- |
| 13C-15N-Ala [M+H]<sup>+</sup> | 1.43 | 94 | 47 | 63 | 8 |
| 13C-15N-Arg [M+H]<sup>+</sup> | 1.36 | 185 | 75 | 63 | 28 |
| 13C-15N-Asn [M+H]<sup>+</sup> | 1.42 | 139 | 77 | 63 | 12 |
| 13C-15N-Asp [M+H]<sup>+</sup> | 1.42 | 139 | 77 | 63 | 12 |
| 13C-15N-Cys [M+H]<sup>+</sup> | 1.38 | 126 | 79 | 63 | 8 |
| 13C-15N-Cystine [M+H]<sup>+</sup> | 1.38 | 249 | 156 | 63 | 8 |
|  | 1.38 | 249 | 77 | 63 | 28 |
| 13C-15N-Gln [M+H]<sup>+</sup> | 1.46 | 154 | 136 | 63 | 4 |
| 13C-15N-Glu [M+H]<sup>+</sup> | 1.47 | 154 | 107 | 63 | 8 |
| 13C-15N-Gly [M+H]<sup>+</sup> | 1.40 | 79 | 50 | 63 | 4 |
|  | 1.40 | 79 | 33 | 63 | 4 |
| 13C-15N-His [M+H]<sup>+</sup> | 1.32 | 165 | 118 | 63 | 12 |
| 13C-15N-Ile [M+H]<sup>+</sup> | 2.77 | 139 | 92 | 63 | 8 |
| 13C-15N-Leu [M+H]<sup>+</sup> | 3.04 | 139 | 92 | 63 | 8 |
| 13C-15N-Lys [M+H]<sup>+</sup> | 1.31 | 155 | 90 | 63 | 16 |
| 13C-15N-Met [M+H]<sup>+</sup> | 2.18 | 156 | 109 | 63 | 8 |
| 13C-15N-Phe [M+H]<sup>+</sup> | 3.50 | 176 | 129 | 63 | 12 |
| 13C-15N-Pro [M+H]<sup>+</sup> | 1.56 | 122 | 75 | 63 | 16 |
| 13C-15N-Ser [M+H]<sup>+</sup> | 1.40 | 110 | 63 | 63 | 8 |
| 13C-15N-Thr [M+H]<sup>+</sup> | 1.44 | 125 | 78 | 63 | 8 |
| 13C-15N-Trp [M+H]<sup>+</sup> | 3.73 | 218 | 200 | 63 | 4 |
|  | 3.73 | 218 | 156 | 63 | 8 |
|  | 3.73 | 218 | 127 | 63 | 28 |
| 13C-15N-Tyr [M+H]<sup>+</sup> | 2.54 | 192 | 145 | 63 | 12 |
| 13C-15N-Val [M+H]<sup>+</sup> | 1.77 | 124 | 77 | 63 | 8 |
| Alanine [M+H]<sup>+</sup> | 1.43 | 90 | 44 | 63 | 8 |
| Arginine [M+H]<sup>+</sup> | 1.36 | 175 | 70 | 63 | 28 |
| Arginine [M+H]<sup>+</sup> | 1.36 | 175 | 60 | 63 | 12 |
| Asparagine [M+H]<sup>+</sup> | 1.42 | 133 | 74 | 63 | 12 |
| Aspartate [M+H]<sup>+</sup> | 1.42 | 134 | 74 | 63 | 12 |
| Cysteine [M+H]<sup>+</sup> | 1.38 | 122 | 76 | 63 | 8 |
|  | 1.38 | 122 | 59 | 63 | 20 |
| Cystine [M+H]<sup>+</sup> | 1.38 | 241 | 152 | 63 | 8 |
|  | 1.38 | 241 | 74 | 63 | 28 |
| Glutamate [M+H]<sup>+</sup> | 1.47 | 148 | 102 | 63 | 8 |
|  | 1.47 | 148 | 84 | 63 | 12 |
|  | 1.47 | 148 | 56 | 63 | 32 |
|  | 1.47 | 147 | 130 | 63 | 4 |
| Glycine [M+H]<sup>+</sup> | 1.40 | 76 | 48 | 63 | 4 |
|  | 1.40 | 76 | 31 | 63 | 4 |
| Histidine [M+H]<sup>+</sup> | 1.32 | 156 | 110 | 63 | 12 |
|  | 1.32 | 156 | 93 | 63 | 24 |
| Isoleucine [M+H]<sup>+</sup> | 2.77 | 132 | 86 | 63 | 8 |
| Leucine [M+H]<sup>+</sup> | 3.04 | 132 | 86 | 63 | 8 |
| Lysine [M+H]<sup>+</sup> | 1.31 | 147 | 84 | 63 | 16 |
| Methionine [M+H]<sup>+</sup> | 2.18 | 150 | 104 | 63 | 8 |
|  | 2.18 | 150 | 61 | 63 | 20 |
|  | 2.18 | 150 | 56 | 63 | 12 |
| Phenylalanine [M+H]<sup>+</sup> | 3.50 | 166 | 120 | 63 | 12 |
|  | 3.50 | 166 | 103 | 63 | 28 |
| Proline [M+H]<sup>+</sup> | 1.56 | 116.1 | 70 | 63 | 16 |
|  | 1.56 | 116 | 43 | 63 | 0 |
| Serine [M+H]<sup>+</sup> | 1.40 | 106 | 60 | 63 | 8 |
|  | 1.40 | 106 | 42 | 63 | 24 |
| Threonine [M+H]<sup>+</sup> | 1.44 | 120 | 74 | 63 | 8 |
|  | 1.44 | 120 | 56 | 63 | 0 |
| Tryptophane [M+H]<sup>+</sup> | 3.73 | 205 | 188 | 63 | 4 |
|  | 3.73 | 205 | 146 | 63 | 8 |
|  | 3.73 | 205 | 118 | 63 | 28 |
| Tyrosine [M+H]<sup>+</sup> | 2.54 | 182 | 136 | 63 | 12 |
|  | 2.54 | 182 | 123 | 63 | 0 |
| Valine [M+H]<sup>+</sup> | 1.77 | 118 | 72 | 63 | 8 |
|  | 1.77 | 118 | 55 | 63 | 0 |

